# Connectomic Alterations Linked to Transdiagnostic Risk for Psychopathology at the Transition to Adolescence

**DOI:** 10.1101/2020.08.21.260927

**Authors:** Chandra Sripada, Mike Angstadt, Saige Rutherford, Aman Taxali, Tristan Greathouse, D. Angus Clark, Luke Hyde, Alex Weigard, Sarah Brislin, Brian Hicks, Mary Heitzeg

**Author notes:** Correspondence: Chandra Sripada, 4250 Plymouth Road, Ann Arbor MI 48109.

## Abstract

**BACKGROUND:** Convergent research identifies a general factor (“P factor”) that confers transdiagnostic risk for psychopathology. However, brain functional connectivity patterns that underpin the P factor remain poorly understood, especially at the transition to adolescence when many serious mental disorders have their onset.

**OBJECTIVE:** Identify a distributed connectome-wide neurosignature of the P factor and assess the generalizability of this neurosignature in held out samples.

**DESIGN, SETTING, AND PARTICIPANTS:** This study used data from the full baseline wave of the Adolescent Brain and Cognitive Development (ABCD) national consortium study, a prospective, population-based study of 11,875 9- and 10-year olds. Data for this study were collected from September 1, 2016 to November 15, 2018 at 21 research sites across the United States.

**MAIN OUTCOMES AND MEASURES:** We produced whole brain functional connectomes for 5,880 youth with high quality resting state scans. We then constructed a low rank basis set of 250 components that captures interindividual connectomic differences. Multi-level regression modeling was used to link these components to the P factor, and leave-one-site-out cross-validation was used to assess generalizability of P factor neurosignatures to held out subjects across 19 ABCD sites.

**RESULTS:** The set of 250 connectomic components was highly statistically significantly related to the P factor, over and above nuisance covariates alone (ANOVA nested model comparison, incremental R-squared 6.05%, χ^2^(250) =412.1, *p*<4.6×10^−10^). In addition, two individual connectomic components were statistically significantly related to the P factor after Bonferroni correction for multiple comparisons (t(5511)= 4.8, *p*<1.4×10^−06^; t(5121)= 3.9, p<9.7×10^−05^). Functional connections linking control networks and default mode network were prominent in the P factor neurosignature. In leave-one-site-out cross-validation, the P factor neurosignature generalized to held out subjects (average correlation between actual and predicted P factor scores across 19 held out sites=0.13; *p*_PERMUTATION_<0.0001). Additionally, results remained significant after a number of robustness checks.

**CONCLUSIONS AND RELEVANCE:** The general factor of psychopathology is associated with connectomic alterations involving control networks and default mode network. Brain imaging combined with network neuroscience can identify distributed and generalizable signatures of transdiagnostic risk for psychopathology during emerging adolescence.

## 1. Introduction

Recent investigations into patterns of covariance across psychiatric symptoms consistently find a general factor of psychopathology, termed the “P factor”, that is associated with most prevalent psychiatric symptoms^1–4^. Concurrently, categorical diagnostic approaches that currently predominate encounter serious issues. Tellingly, one persistent problem has been excess overlap across disorders in symptoms^5,6^, neural mechanisms^7,8^, and genetic risk factors^9,10^—a problem that the could be readily explained if a domain general P factor drives cooccurrence of symptoms irrespective of diagnostic boundaries. Despite these compelling features of the P factor model, key gaps in knowledge remain, especially regarding the neural mechanisms that produce broad liabilities to diverse psychopathologies and the developmental pathways through which these broad liabilities operate.

Network neuroscience^11–14^ is well positioned to help fill in this gap in knowledge. The human brain is organized into a number of large-scale connectivity networks^15,16^, and there is growing understanding of distinct information processing functions implemented by these networks and by interacting network ensembles. Recent psychological models of the P factor emphasize executive dysfunction and impulsivity^3,17,18^, suggesting the possibility that deficits in control networks (e.g., fronto-parietal network, dorsal attention network)^19,20^, involved in attention and cognitive control, are implicated in producing the P factor—a possibility that awaits detailed investigation.

Network neuroscience can also illuminate the developmental pathways that lead to psychopathology^21–23^. Brain networks undergo massive maturation during adolescence^24–26^. Importantly, this is also the time that many serious mental disorders first emerge^5,21,27^. A critical goal for psychiatric neuroscience is to develop brain-based objective markers of risk that can identify vulnerable youth who can most benefit from targeted interventions^21^. Because of the breadth of the P factor’s posited effects on diverse forms of psychiatric symptomatology, identification of network alterations during youth that are associated with the P factor is particularly pressing.

The current study examines connectome-wide functional connectivity patterns associated with the P factor in a sample of 11,875 9 and 10-year olds in the Adolescent Brain and Cognitive Development (ABCD) national consortium study, Release 2.1^28,29^. Recently, our group constructed and validated a P factor model^30^ in ABCD from the Child Behavior Checklist^31^ (CBCL)-parent report using bifactor modeling. For the present study, we produced resting state connectomes for 5,880 youth who met stringent neuroimaging quality control standards. We next applied a multivariate approach, brain basis set (BBS)^32–34^, to link whole-brain functional connectivity patterns to the P factor. BBS takes advantage of the fact that, although functional connectomes are large and complex, encompassing tens of thousands of connections, there is massive redundancy in the set of connections that differ across people. This allows a small set of components—we used 250 in the present study—to capture most meaningful interindividual variation^32,33^. To establish multivariate relationships between this basis set of connectomic components and the P factor, we adopted a two-pronged analysis approach that leverages standard parametric statistical methods (well suited for complex multilevel designs with covariates and sample weighting) and newer cross-validation approaches (well suited for assessing generalizability to unseen data). Both analyses convergently demonstrate that the P factor is associated with brain-wide altered connectivity patterns prominently implicating connections within and between control networks and processing networks.

## 2. Methods

### 2.1. Sample and Data

The ABCD study is a multisite longitudinal study with 11,875 children between 9-10 years of age from 21 sites across the United States. The study conforms to the rules and procedures of each site’s Institutional Review Board, and all participants provide informed consent (parents) or assent (children). Detailed description of recruitment procedures^35^, assessments^36^, and imaging protocols^37^ are available elsewhere. The ABCD data used in this report came from NDA Study 721,10.15154/1504041, which can be found at https://nda.nih.gov/study.html?id=721.

### 2.2. Data Acquisition, fMRI Preprocessing, and Connectome Generation

Imaging protocols were harmonized across sites and scanners. High spatial (2.4 mm isotropic) and temporal resolution (TR=800 ms) resting state fMRI was acquired in four separate runs (5min per run, 20 minutes total, full details are described in ^38^). The entire data pipeline described below was run through automated scripts on the University of Michigan’s high-performance cluster, and is described below, with additional detailed methods automatically generated by fRMIPrep software provided in the Supplement. Code for running the analyses can be found at https://github.com/SripadaLab/ABCD_Resting_Psychopathology.

Preprocessing was performed using fMRIPrep version 1.5.0^39^, a Nipype^40^ based tool. Full details of the fMRIPrep analysis can be found in supplemental materials. Briefly, T1-weighted (T1w) and T2-weighted images were run through recon-all using FreeSurfer v6.0.1. T1w images were also spatially normalized nonlinearly to MNI152NLin6Asym space using ANTs 2.2.0. Each functional run was corrected for fieldmap distortions, rigidly coregistered to the T1, motion corrected, and normalized to standard space. ICA-AROMA was run to generate aggressive noise regressors. Anatomical CompCor was run and the top 5 principal components of both CSF and white matter were retained. Functional data were transformed to CIFTI space using the method provided on the HCP Wiki (https://wiki.humanconnectome.org/display/PublicData/HCP+Users+FAQ#HCPUsersFAC-9.HowdolmapdatabetweenFreeSurferandHCP?). All preprocessed data were visually inspected at two separate stages to ensure only high-quality data was included: After co-registration of the functional data to the structural data and after registration of the functional data to MNI template space.

Connectomes were generated for each functional run using the Gordon 333 parcel atlas^41^, augmented with parcels from high-resolution subcortical^42^ and cerebellar^43^ atlases. Volumes exceeding a framewise displacement threshold of 0.5mm were marked to be censored. Covariates were regressed out of the time series in a single step^44^, including: linear trend, 24 motion parameters (original translations/rotations + derivatives + quadratics, aCompCorr 5 CSF and 5 WM components and ICA-AROMA aggressive components, high pass filtering at 0.008Hz, and censored volumes. Next, correlation matrices were calculated for each run. Each matrix was then Fisher r-to-z transformed, and then averaged across runs for each subject yielding their final connectome.

### 2.3. Constructing Structural Model of Psychopathology

The general psychopathology factor (P-factor) used here is based on the parent-rated Child Behavior Checklist (CBCL; age 6 to 18 form^31^). A bifactor model was fit to eight CBCL scales, with a general P factor that all scales loaded onto (average scale loading = .69) and two specific factors. This model is described in detail in the Supplement as well our previous studies, where we compared this model to 14 alternatives^30^ (finding consistent P factors across diverse model specifications, rs>0.90) and we delineated nomological networks of the P factor across diverse ABCD variables^45^.

### 2.4. Inclusion/Exclusion

There are 11,875 subjects in the ABCD Release 2.0.1 dataset. Screening was initially done using ABCD raw QC to limit to subjects with 2 or more good runs of resting data as well as a good T1 and T2 image (QC score, protocol compliance score, and complete all =1). This resulted in 9580 subjects with 2 or more runs that entered preprocessing. Each run was subsequently visually inspected for registration and warping quality, and only those subjects who still had 2 or more good runs were retained (N=8858). After connectome generation, runs were excluded if they had less than 4 minutes of uncensored data, and next subjects were retained only if they had to 2 or more good runs (N=6568). Finally, subjects who were missing data required for regression modeling (P factor scores or any of the nuisance covariates) were dropped, and then sites with fewer than 75 subjects were dropped. This left us with N=5880 subjects across 19 sites for the whole sample multiple regression analysis and leave-one-site-out cross-validation.

### 2.5. Constructing a Brain Basis Set (BBS)

BBS is a validated multivariate method that uses principal components dimensionality reduction to produce a basis set of components that are then associated with phenotypes^46,32^. We select the top 250 components for our basis set based on previous work showing that 50-100 components per 1000 subjects captures most meaningful variance without overfitting^32^.

### 2.6. Whole Sample Multiple Regression Analysis

To assess the multivariate relationship between the 250-component brain basis set and the P factor, we used multi-level multiple regression modeling, with the P factor as outcome variable and expression scores for the 250 components as predictors. Gender, race, age, mean FD, and mean FD squared were entered as fixed effect covariates, with family id and ABCD site entered as random effects (family nested within site).

To help convey overall patterns across the entire multiple regression model, we constructed a “consensus” component map. We multiplied each connectomic component with its associated beta from the preceding multiple regression model. Next, we summed across all 250 components yielding a single map, and thresholded the entries at z=2.

### 2.7. Leave-One-Site-Out Cross Validation

To assess generalizability of BBS-based regression models, we used leave-one-site-out crossvalidation. In each fold of the cross-validation, data from one of the 19 sites served as the held-out test dataset and data from the other 18 sites served as the training dataset. Additionally, to ensure separation of train and test datasets, at each fold of the cross-validation, a new PCA was performed on the training dataset yielding a 250-component basis set.

### 2.8. Accounting for Covariates in Cross-Validation Framework

In each fold of cross-validation, BBS models were trained in the train partition with the following covariates: gender, race, age, age squared, mean FD and mean FD squared. To maintain strict separation between training and test datasets, regression coefficients for the covariates learned from the training sample are applied to the test sample, and the variance they explain is subtracted away. This procedure, described in detail in our previous publication^34^, yields an estimate of the contribution of brain components alone in predicting test subject P factor scores, excluding the contribution of the nuisance covariates. Note that by employing leave-one-site-out, members of twinships and sibships are never present in both training and test samples.

### 2.9. Permutation Testing

We assessed the significance of all cross-validation-based correlations with non-parametric permutation tests in which we randomly permuted the 5,880 subjects’ P factor scores 10,000 times, as described in detail in the Supplement.

## 3. Results

### 3.1. A Distributed Pattern of Network Alterations Is Highly Statistically Significantly Related to the P Factor

We used multiple regression analysis to quantify the relationship between brain connectivity patterns and the P factor. In a multi-level regression model, we entered 250 brain components as predictors and the P factor as outcome variable, with race, gender, head motion, ABCD site, and family id as nuisance covariates. We found that compared to a covariates-only model, the 250 brain components yielded an 6.05% increase in variance explained, a highly statistically significant increase (χ^2^(250) = 412.1, *p* < 4.6×10^−10^; Figure 1). We in addition created a consensus connectome, an importance-weighted composite of components, that represents a multivariate neurosignature of the P factor (Figure 2). Two motifs are particularly prominent in the neurosignature: 1) altered connectivity within the 14 large-scale networks, especially in DMN and sensory networks; and 2) altered connectivity within control networks and between control networks and DMN. In particular, connections within large-scale networks (along the diagonal of the figure) represent 8.48% of the connectome, but 23.65% of the suprathreshold connections in the neurosignature. Additionally, cross-network connections involving control networks (FPN, CO, DAN, VAN) and DMN represent 11.58% of the connectome, but 31.83% of the suprathreshold connections in the neurosignature.

**Figure 1.**
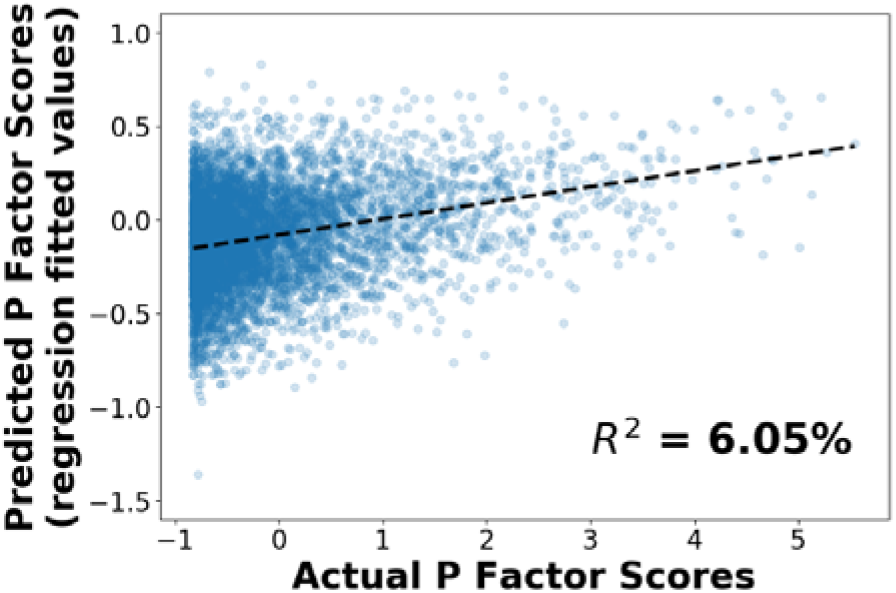
Fitted Relationship Between 250 Connectivity Components and the P Factor in 5,880 Youths. We conducted a multi-level regression with 250 connectivity components as predictors and the P Factor as outcome variable, controlling for demographic covariates and clustering due to family and site. The scatter plot reflects the fitted fixed effect relationship between predictors and the P factor (incremental R squared of 250 components over and above a covariates-only model is 6.05%).

**Figure 2:**
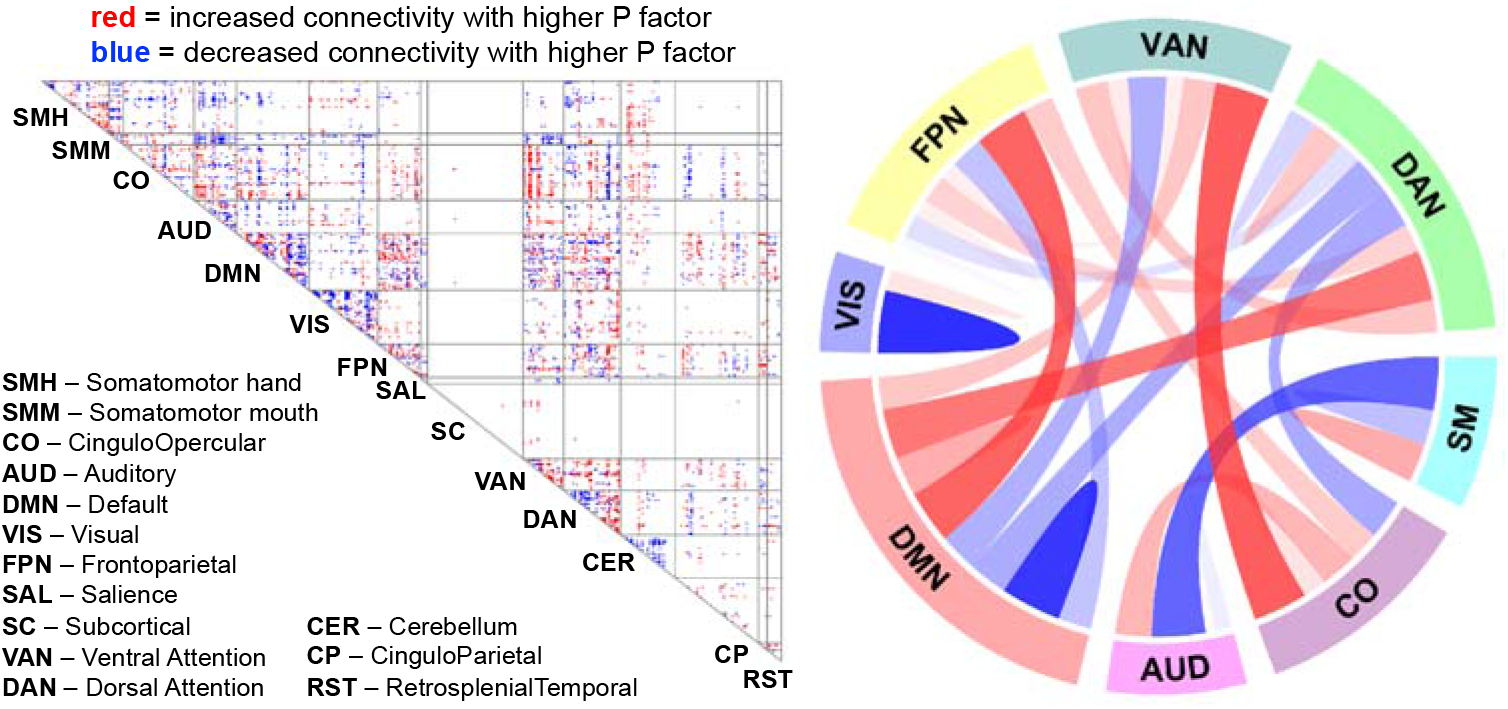
Multivariate Connectomic Neurosignature of the P Factor. Using multi-level regression modeling, we observed a highly statistically significant relationship between an individual differences basis set of 250 connectivity components and the P factor. Based on the fitted model coefficients, we created a consensus connectome, an importance-weighted composite of components, that represents a multivariate neurosignature of the P factor. (Left Panel) Connection-level map of the consensus connectome. This map shows altered within-network connectivity (triangles along the diagonal of the left panel) at several networks including DMN. It also shows altered connectivity involving control networks (FPN, CO, VAN, DAN) and DMN. (Right Panel) Network-level map of the consensus connectome, in which width of arcs reflects number of suprathreshold connections. Note, to enhance interpretability, some networks are not shown (see Supplement).

### 3.2. Two Connectivity Components are Statistically Significantly Related to the P factor

We next examined statistical significance of individual connectivity components within the multiple regression model, using Bonferroni correction to control family-wise error while making 250 statistical comparisons. After this correction, we found that two components remained statically significantly associated with the P factor (t(5511)= 4.8, p<1.4×10^−06^; t(5121)= 3.9, p<9.7×10^−05^). Connectivity patterns from these components are shown in Figure 3 (see its associated figure caption for descriptions of these findings). Importantly, there were 27 components that were significantly associated the P factor at p<0.05, uncorrected (where only 12.5 are expected by chance), indicating that the P factor is associated with numerous relatively subtle connectome-wide alterations.

**Figure 3:**
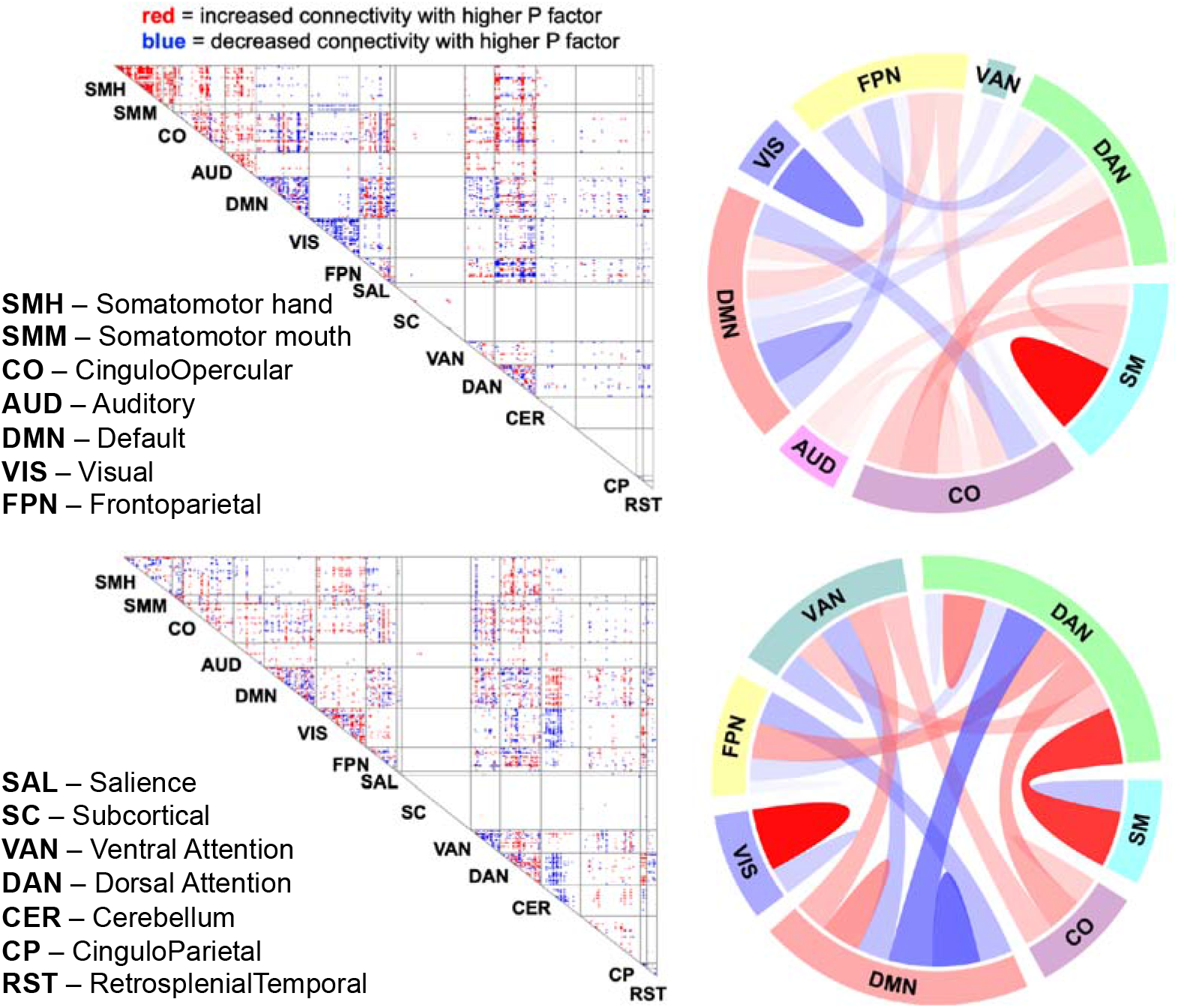
Connectivity Components Associated with the P Factor. Two individual connectivity components were statistically significantly related to the P factor (after Bonferroni correction for 250 comparisons). Both components involve alterations in within-network connectivity, especially in SMH, DMN, VN, and DAN, as well as altered connectivity involving control networks and DMN. (Left side) Connection-level map showing suprathreshold connections. (Right Side) Network-level map, in which width of arcs reflects number of suprathreshold connections. Note, to enhance interpretability, some networks are not shown (see Supplement).

### 3.3. Multivariate Neurosignatures of the P factor Generalize to Held Out Samples

We next examined the relationship between these 250 components and the P factor using a predictive modeling framework with leave-one-site-out cross-validation. A regression model was trained for connectome-based prediction of the P factor at all sites except one, this model was used to derive predictions of the P factor at the held out site, and this process was repeated with each site held out. To maintain separation between training and testing samples, we learned new connectivity components and covariate regression coefficients in each training sample and applied them to the testing sample. The correlation between actual and predicted P factor scores was 0.13, a value higher than all 10,000 values in a permutation-based null distribution (p_PERM_<0.0001; Figure 4).

**Figure 4.**
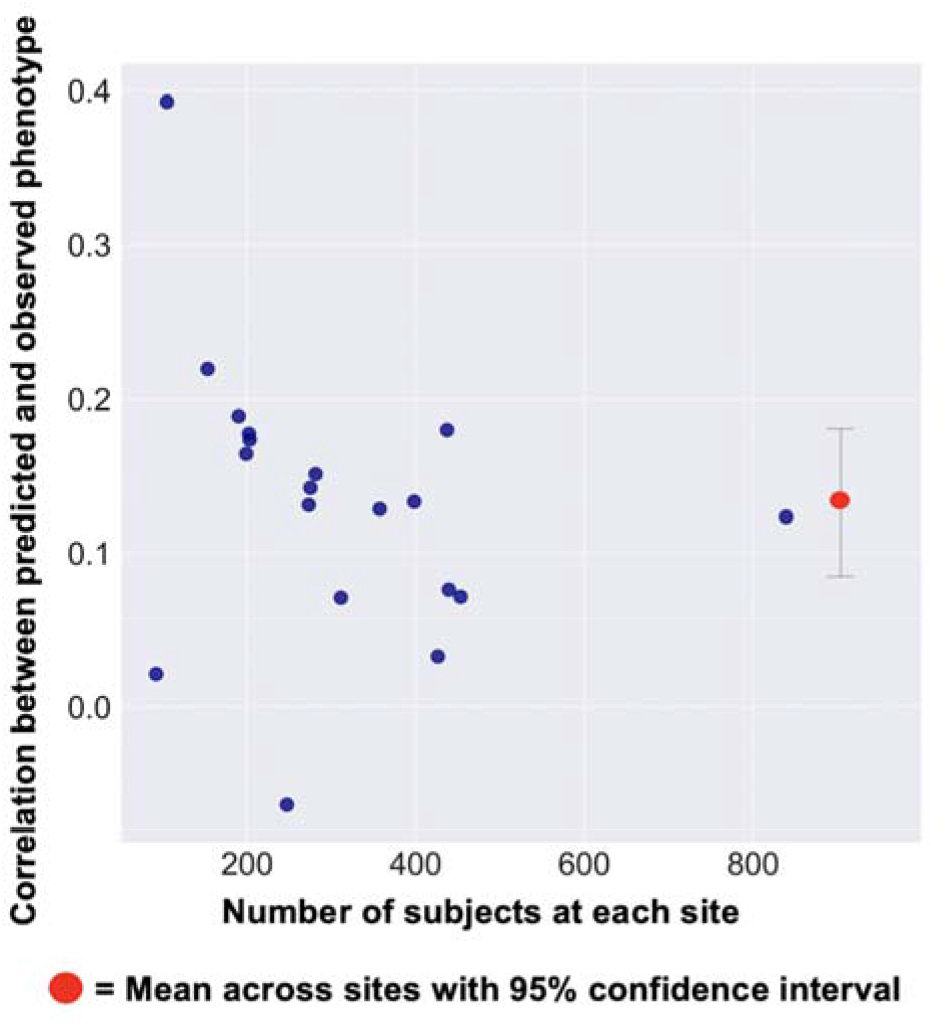
Site-Specific Correlations Between Predicted and Actual P Factor Scores in Leave-One-Site-Out Cross-Validation Analysis.

### 3.4. Multivariate Neurosignatures Remain Significantly Related to the P factor Across a Number of Tests of Robustness

We next assessed the robustness of our multiple regression analysis and leave-one-site-out cross-validation analysis by changing key elements of the analysis streams. We tested models that utilized: 1) Log-transformed P factor scores to improve their distributional properties; 2) Additional ABCD demographic covariates (household income, highest parental education, household marital status); 3) An ultra-low head motion sample (mean FD<0.2; *N* = 2,721); 4) P factor models learned in the training sample and applied to the held out sample (to create total separation of train and test partitions); 5) Sampling weights, based on American Community Survey proportions^47^, which make findings more generalizable by approximating a representative population. As shown in Table 1, all models remained highly statistically significant across all these analyses, confirming the robustness of our analysis.

**Table 1:**
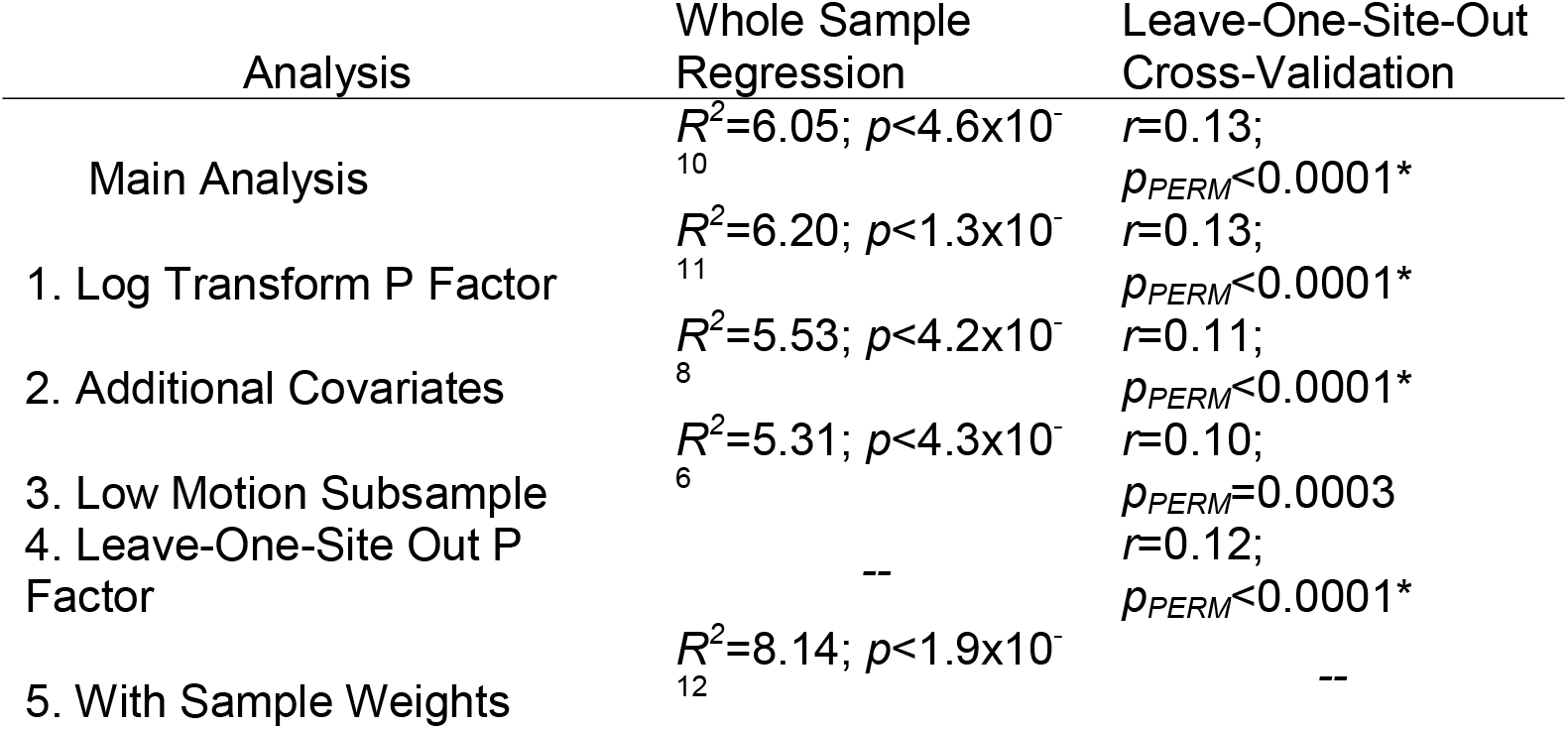
Summary of Additional Analyses to Assess Robustness. We assessed the sensitivity of our main analyses (top row) to modeling choices with a number of robustness checks (rows labeled 1-5). We found results remained statistically significant across all these analyses. * = observed correlation was higher than all 10,000 correlations in the permutation distribution.

## 4. Discussion

This study investigated brain-wide connectomic alterations associated with the general factor of psychopathology (“P factor”), a broad liability for diverse forms of psychopathology, in 5,880 9- and 10-year olds in the ABCD multi-site sample. By combining large sample size, reliable multivariate methods, cross-validation to assess generalizability, sampling weights to approximate representativeness, and a number of additional tests of model robustness, we provide the strongest evidence yet that the P factor is associated with distributed changes in connectivity patterns of the brain’s intrinsic functional architecture. We found especially prominent alterations in connections within large-scale networks and connections linking control networks and DMN. Overall, our results highlight the potential for neuroimaging to detect and delineate network abnormalities associated with broad liabilities for psychopathology during the transition to adolescence, a key developmental window in which many serious psychopathologies first emerge.

Previous studies tended to use case control designs to characterize brain network abnormalities associated with individual disorders^48^, where these abnormalities were assumed to reflect disorder-specific pathophysiology. Accumulated results, however, suggest observed network alterations often lack specificity. For example, two prominent network motifs we observed in the present study, reduced connectivity within DMN and altered control networks/DMN connectivity, have elsewhere been demonstrated in a number of individual psychiatric disorders, including hypoconnectivity of DMN in autism^49^, schizophrenia^50^, and ADHD^46,51,52^, and reduced DMN/TPN anti-correlation in schizophrenia^53–55^, bipolar disorder^56^, and ADHD^46,52^. The present study potentially explains this lack of specificity by linking motifs such as these instead to the P factor. The P factor represents a broad liability to diverse forms of psychopathology, and thus P-linked connectomic motifs would be expected to show up non-specificaIly across diverse case-control disorder comparisons.

It is noteworthy that these same two connectivity motifs, i.e., connectivity within DMN and control networks/DMN connectivity, are known to undergo intense maturation during childhood and adolescence^26^. In particular, during this developmental period, links within DMN exhibit massive increased connectivity^57,46^ and links involving control networks and DMN exhibit substantial reconfiguration^57–59,46^, with a predominant pattern of segregation (increasing anti-correlation)^58,46^. These observations are suggestive of graphical colocation of P-effects and maturation-effects in the connectome, with P associated with a more immature connectivity pattern. Future studies should quantitatively examine this hypothesis, by using formal tests for graphical colocation in cross-sectional data^59^ or by quantifying P-associated maturation patterns in future waves of ABCD longitudinal data. Studies such as these could provide direct evidence that the P factor reflects neurodevelopmental dysmaturation, inviting further research into how to mitigate “miswired” connectivity patterns^60^ or prospectively block their emergence.

It is interesting to interpret our results in light of recent proposals for the psychological basis of the P factor. We recently examined^45^ the nomological networks of the P factor in the ABCD sample (n=11,875), finding that P is associated with three psychological variables: higher levels of fear/distress emotions, higher impulsivity, and reduced neurocognition. This aligns with proposals from other researchers^3,17,18^ that P represents heightened generation of impulses (both negative fear/distress emotions as well as reward-seeking impulses) and reduced executive functioning. Consistent with this idea, our connectomic results highlight altered connectivity involving control networks (FPN, CO, DAN, VAN) and processing networks (DMN, visual network, auditory network). Notably other recent functional connectivity studies of the P factor also found alterations in processing networks, including visual network^61^ and somatomotor network^62^. Control networks are involved in working memory and attention and are sources of cognitive control signals^19,20^, while processing networks are involved in stimulus processing and spontaneous thought^63^ and are targets of cognitive control^64,65^. Thus, known functional profiles of networks found here to be associated with the P factor align well with current models of the psychological factors that contribute to higher levels of P.

While the brain/behavior associations identified in this study are statistically reliable and generalize to held out samples, they are nonetheless modest in size, and several points are relevant. First, we systematically assessed the ability of the multivariate neurosignature of the P factor to generalize to brain imaging data from 19 held out sites, each employing their own fMRI scanners. Few studies of psychopathology seek to systematically assess cross-site generalizability in this way (cf.^33^). Direct comparison of our multivariate cross-validation effect size with within-sample effect sizes from other studies would thus be inappropriate. Such comparison would be especially inapt if the comparison study used more standard mass univariate analysis methods (rather than multivariate methods), because conducting thousands of statistical tests produces inflated p-values (which is widely known) as well as inflated effect sizes^66^ (which is less well appreciated). Second, this study was conducted in 9- and 10-year old youth, many of whom had relatively low levels of psychiatric symptomatology. It is expected that subjects’ psychopathology load will rise during the course of adolescence^27^, and it is possible that brain/behavior relationships will correspondingly increase, a hypothesis that can be directly tested in future waves of longitudinal ABCD data. Third, this study exclusively examined brain/behavior relationships with resting state functional connectomes. Other lines of work link the P factor to structural alterations^67^ (gray matter thinning) and white matter tract changes^68,69^. Thus, it is possible that multi-modal fusion methods that aggregate information across modalities will further boost the size of brain/behavior relationships.

In sum, using a rigorous multivariate approach combined with cross-validation in held out samples, we established that during emerging adolescence, the P factor is associated with distributed alterations in control networks and DMN, critical elements of the brain’s intrinsic functional architecture. These findings set the stage for future studies in the ABCD sample, leveraging longitudinal waves of data to trace the psychological and neural progression of the P factor during a critical window of vulnerability to mental illness that spans adolescence to young adulthood.

## Acknowledgements

Data used in the preparation of this article were obtained from the Adolescent Brain Cognitive Development (ABCD) Study (https://abcdstudy.org), held in the NIMH Data Archive (NDA). This is a multisite, longitudinal study designed to recruit more than 10,000 children age 9-10 and follow them over 10 years into early adulthood. The ABCD Study is supported by the National Institutes of Health and additional federal partners under award numbers U01DA041022, U01DA041028, U01DA041048, U01DA041089, U01DA041106, U01DA041117, U01DA041120, U01DA041134, U01DA041148, U01DA041156, U01DA041174, U24DA041123, and U24DA041147. A full list of supporters is available at https://abcdstudy.org/nih-collaborators. A listing of participating sites and a complete listing of the study investigators can be found at https://abcdstudy.org/principal-investigators.html. ABCD consortium investigators designed and implemented the study and/or provided data but did not necessarily participate in analysis or writing of this report. This manuscript reflects the views of the authors and may not reflect the opinions or views of the NIH or ABCD consortium investigators. The ABCD data repository grows and changes over time. The ABCD data used in this report came from NDA Study 721, 10.15154/1504041, which can be found at https://nda.nih.gov/study.html?id=721.

This work was supported by the following grants from the United States National Institutes of Health, the National Institute on Drug Abuse, and the National Institute on Alcohol Abuse and Alcoholism: R01MH107741 (CS), U01DA041106 (CS, LH, MH, BH), T32 AA007477 (DC, AW). In addition, CS was supported by a grant from the Dana Foundation David Mahoney Neuroimaging Program. This research was supported in part through computational resources and services provided by Advanced Research Computing at the University of Michigan, Ann Arbor.

## Supplement

### Quality Control-Resting State Functional Connectivity Plot

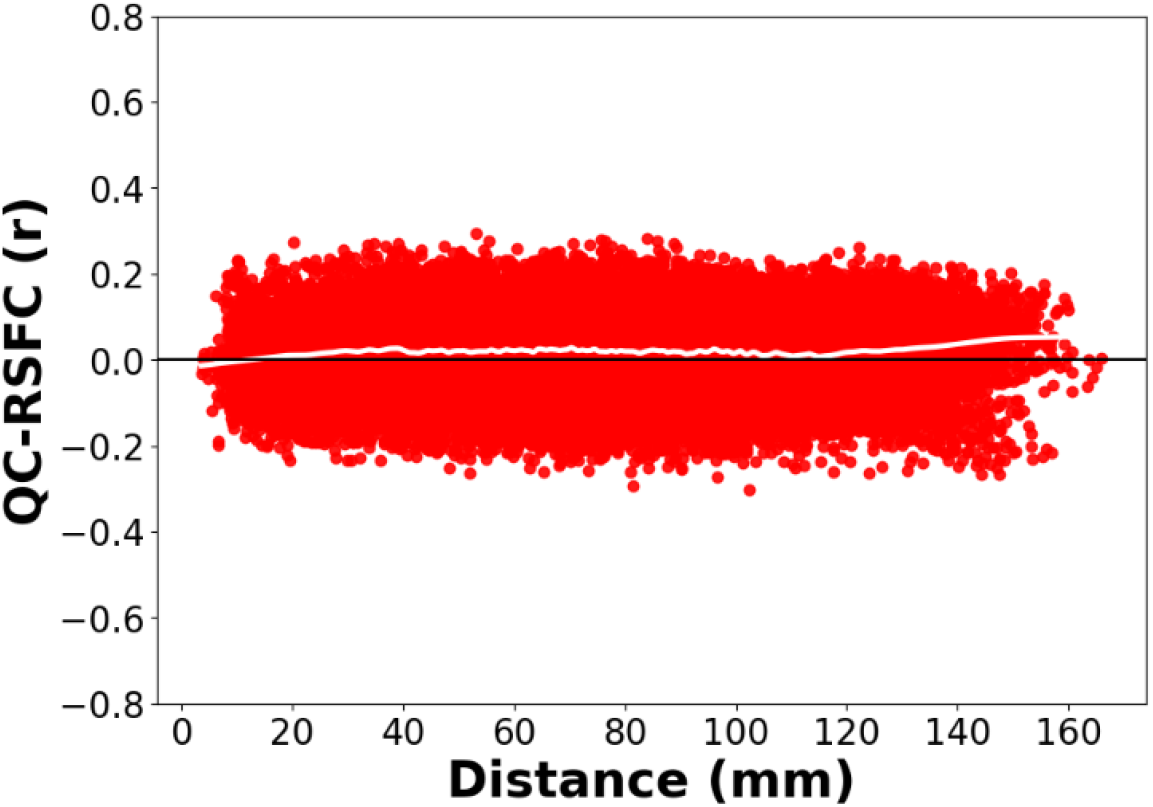

We used multiple procedures to control the effect of head motion on brain-behavior relationships, which are listed in §2.2 in the Main Manuscript. To assess the effectiveness of these procedures, we produced a quality control resting state functional connectivity (QC-RSFC) plot^1,2^. This plot shows the relationship between mean framewise displacement and connectivity for edges binned by distance. Motion effects produce a sloped line (distance-dependent artifact), while a flat line is indicative of minimal motion-related effects. The RSFC-QC plot for our ABCD resting state data showed a flat line, providing additional evidence that our stringent motion correction strategies were effective.

### P Factor Modeling

A general P-factor with two orthogonal specific factors were modeled using the parent-rated Child Behavior Checklist (CBCL; age 6 to 18 form^3^). We fit a bifactor model to the eight CBCL scales (Withdrawn, Somatic Complaints, Anxious/Depressed, Social Problems, Thought Problems, Attention Problems, Delinquent Behavior, and Aggressive Behavior). In this model there was a general P factor that all scales loaded onto (average scale loading on P = .69), and internalizing and externalizing specific factors (average scale loading = .43). This model fit well based on conventional fit thresholds (χ^2^ = 747.73, df = 16, *p* < .001; RMSEA = .062; CFI = .985; TLI = .974; SRMR = .015) and was chosen for its good model fit and theoretical interpretability. Importantly, we elsewhere demonstrate that across a variety of alternative specifications of hierarchical models of psychopathology in ABCD, the resulting P factors that emerge yield highly similar rank ordering of subjects (rs>0.90)^4^, rendering decisions about which specific modeling strategy to adopt less consequential.

### Network Visualizations

We created circular network visualizations of whole brain connectomes (see Figures 2 and 3 in the Main Manuscript) using the circlize software library in R. To increase interpretability, SMH and SMM were combined into a single somatomotor network (SM) and five networks that were not extensively implicated were omitted (SAL, SC, CER, CP, RST). In addition, cells in which suprathreshold connections were less than 7.5% of the total possible connections were omitted.

### Permutation Framework

The distribution under chance of correlations between BBS-based predictions of neurocognitive scores and observed neurocognitive scores was generated by randomly permuting the 5,880 subjects’ P factor scores 10,000 times. At each iteration, we performed the leave-one-site out cross validation procedure described in the main manuscript, including refitting BBS models at each fold of the cross-validation. We then recalculated the average correlation across folds between predicted versus actual P factor scores. The average correlation across folds that was actually observed was located in this null distribution in terms of rank, and statistical significance was set as this rank value divided by 10,000.

Since the BBS models fit at each iteration of the permutation test included covariates, the procedure of Freedman and Lane^5^ was followed. In brief, a BBS model was first estimated with nuisance covariates alone, residuals were formed and were permuted. The covariate effect of interest was then included in the subsequent model, creating an approximate realization of data under the null hypothesis, and the statistical test of interest was calculated on this data (see FSL Randomise http://fsl.fmrib.ox.ac.uk/fsl/fslwiki/Randomise/Theory for a neuroimaging implementation).

### Robustness Checks

We performed a number of additional analyses to assess the robustness of our results under alternative model specifications. One alternative approach derived P factor scores with strict separation between training and testing partitions. In particular, for the train partition subjects, we fit a bifactor model to CBCL subscales as described in the Main Manuscript (§2.3) and derived P factor scores. We then applied this same fitted model to the held-out test subjects to obtain their P factor scores. This process was repeated for all folds of the leave-one-site-out cross-validation.

**Supplemental Table 1:** Demographic Characteristics of Included Versus Excluded Subjects

|  | Included | Excluded |
| --- | --- | --- |
| N | 5880 | 5995 |
| Age (mean (s.d.)) | 9.97 (0.62) | 9.88 (0.62) |
| Female (%) | 2973 (50.6) | 2708 (45.2) |
| Race Ethnicity (%) |  |  |
| White | 3543 (60.2) | 2581 (43.1) |
| Black | 663 (11.3) | 1095 (18.3) |
| Hispanic | 1036 (17.6) | 1343 (22.4) |
| Asian | 86 (1.5) | 177 (3.0) |
| Other | 552 (9.4) | 642 (10.7) |
| No answer | -- | 157 (2.6) |
| Highest Parental Education (%) |  |  |
| < HS Diploma | 173 (2.9) | 410 (6.8) |
| Bachelor | 1649 (28.0) | 1339 (22.3) |
| HS Diploma/GED | 386 (6.6) | 734 (12.2) |
| Post Graduate Degree | 2223 (37.8) | 1782 (29.7) |
| Some College | 1449 (24.6) | 1590 (26.5) |
| No answer | -- | 140 (2.3) |
| Household Marital Status –<br>Married (%) | 4305 (73.2) | 3604(60.1) |
| Household Income (%) |  |  |
| <50K | 1486 (25.3) | 1692 (28.2) |
| >=100k | 2617 (44.5) | 1941 (31.9) |
| >=50k & <100K | 1777 (30.2) | 1261 (21.0) |
| No answer | -- | 1128 (18.8) |

## Supplemental Methods

### FMRIPrep Methods

The following was generated automatically by fMRIPrep software and is copied here unchanged:

Results included in this manuscript come from preprocessing performed using *fMRIPrep* 1.5.0 (Esteban, Markiewicz, et al. (2018); Esteban, Blair, et al. (2018); RRID:SCR_016216), which is based on *Nipype* 1.2.2 (Gorgolewski et al. (2011); Gorgolewski et al. (2018); RRID:SCR_002502).

#### Anatomical data preprocessing

The T1-weighted (T1w) image was corrected for intensity non-uniformity (INU) with N4BiasFieldCorrection (Tustison et al. 2010), distributed with ANTs 2.2.0 (Avants et al. 2008, RRID:SCR_004757), and used as T1w-reference throughout the workflow. The T1w-reference was then skull-stripped with a *Nipype* implementation of the antsBrai⊓Extraction.sh workflow (from ANTs), using OAS1S3OANTs as target template. Brain tissue segmentation of cerebrospinal fluid (CSF), white-matter (WM) and gray-matter (GM) was performed on the brain-extracted T1w using fast (FSL 5.0.9, RRID:SCR_002823, Zhang, Brady, and Smith 2001). Brain surfaces were reconstructed using recon-all (FreeSurfer 6.0.1, RRID:SCR_001847, Dale, Fischl, and Sereno 1999), and the brain mask estimated previously was refined with a custom variation of the method to reconcile ANTs-derived and FreeSurfer-derived segmentations of the cortical gray-matter of Mindboggle (RRID:SCR_002438, Klein et al. 2017). Volume-based spatial normalization to one standard space (MNI152NLin6Asym) was performed through nonlinear registration with antsRegistration (ANTs 2.2.0), using brain-extracted versions of both T1w reference and the T1w template. The following template was selected for spatial normalization: *FSL’s MNIICBM152 non-linear 6th Generation Asymmetric Average Brain Stereotaxic Registration Model* [Evans et al. (2012), RRID:SCR_002823; TemplateFlow ID: MNI152NLin6Asym].

#### Functional data preprocessing

For each of the 10 BOLD runs found per subject (across all tasks and sessions), the following preprocessing was performed. First, a reference volume and its skull-stripped version were generated using a custom methodology *of fMRIPrep.* A deformation field to correct for susceptibility distortions was estimated based on two echo-planar imaging (EPI) references with opposing phase-encoding directions, using 3dQwarp Cox and Hyde (1997) (AFNI 20160207). Based on the estimated susceptibility distortion, an unwarped BOLD reference was calculated for a more accurate co-registration with the anatomical reference. The BOLD reference was then co-registered to the T1w reference using bbregister (FreeSurfer) which implements boundary-based registration (Greve and Fischl 2009). Co-registration was configured with six degrees of freedom. Head-motion parameters with respect to the BOLD reference (transformation matrices, and six corresponding rotation and translation parameters) are estimated before any spatiotemporal filtering using mcflirt (FSL 5.0.9, Jenkinson et al. 2002). The BOLD timeseries, were resampled to surfaces on the following spaces: *fsaverage5.* The BOLD timeseries (including slice-timing correction when applied) were resampled onto their original, native space by applying a single, composite transform to correct for headmotion and susceptibility distortions. These resampled BOLD time-series will be referred to as *preprocessed BOLD in original space*, or just *preprocessed BOLD.* The BOLD time-series were resampled into standard space, generating a *preprocessed BOLD run in [‘MNI152NLin6Asym’] space.* First, a reference volume and its skull-stripped version were generated using a custom methodology *of fMRIPrep.* Automatic removal of motion artifacts using independent component analysis (ICA-AROMA, Pruim et al. 2015) was performed on the *preprocessed BOLD on MNI space* time-series after removal of nonsteady state volumes and spatial smoothing with an isotropic, Gaussian kernel of 6mm FWHM (full-width half-maximum). Corresponding “non-aggresively” denoised runs were produced after such smoothing. Additionally, the “aggressive” noise-regressors were collected and placed in the corresponding confounds file. Several confounding timeseries were calculated based on the *preprocessed BOLD:* framewise displacement (FD), DVARS and three region-wise global signals. FD and DVARS are calculated for each functional run, both using their implementations in *Nipype* (following the definitions by Power et al. 2014). The three global signals are extracted within the CSF, the WM, and the whole-brain masks. Additionally, a set of physiological regressors were extracted to allow for component-based noise correction *(CompCor*, Behzadi et al. 2007). Principal components are estimated after high-pass filtering the *preprocessed BOLD* time-series (using a discrete cosine filter with 128s cut-off) for the two CompCor variants: temporal (tCompCor) and anatomical (aCompCor). tCompCor components are then calculated from the top 5% variable voxels within a mask covering the subcortical regions. This subcortical mask is obtained by heavily eroding the brain mask, which ensures it does not include cortical GM regions. For aCompCor, components are calculated within the intersection of the aforementioned mask and the union of CSF and WM masks calculated in T1w space, after their projection to the native space of each functional run (using the inverse BOLD-to-T1w transformation). Components are also calculated separately within the WM and CSF masks. For each CompCor decomposition, the *k* components with the largest singular values are retained, such that the retained components’ time series are sufficient to explain 50 percent of variance across the nuisance mask (CSF, WM, combined, or temporal). The remaining components are dropped from consideration. The head-motion estimates calculated in the correction step were also placed within the corresponding confounds file. The confound time series derived from head motion estimates and global signals were expanded with the inclusion of temporal derivatives and quadratic terms for each (Satterthwaite et al. 2013). Frames that exceeded a threshold of 0.5 mm FD or 1.5 standardised DVARS were annotated as motion outliers. All resamplings can be performed with *a single interpolation step* by composing all the pertinent transformations (i.e. head-motion transform matrices, susceptibility distortion correction when available, and co-registrations to anatomical and output spaces). Gridded (volumetric) resamplings were performed using antsApplyTransforms (ANTs), configured with Lanczos interpolation to minimize the smoothing effects of other kernels (Lanczos 1964). Non-gridded (surface) resamplings were performed using mri_vol2surf (FreeSurfer).

Many internal operations of *fMRIPrep* use *Nilearn* 0.5.2 (Abraham et al. 2014, RRID:SCR_001362), mostly within the functional processing workflow. For more details of the pipeline, see the section corresponding to workflows in *fMRIPrep’s* documentation.

### Copyright Waiver

The above boilerplate text was automatically generated by fMRIPrep with the express intention that users should copy and paste this text into their manuscripts *unchanged.* It is released under the CC0 license.

